# Skin microbiome modulation induced by probiotic solutions

**DOI:** 10.1101/423285

**Authors:** Bernhard Paetzold, Jesse R. Willis, Joao Lima, Nastassia Knodlseder, Sven R. Quist, Toni Gabaldón, Marc Güell

## Abstract

**Background:** The skin is colonized by a large number of microorganisms, of which most are beneficial or harmless. However, disease states of skin have specific microbiome compositions that are different from those of healthy skin. Gut microbiome modulation through fecal transplant has proven as a valid therapeutic strategy in diseases such as *Clostridium difficile* infections. Therefore, techniques to modulate the skin microbiome composition may become an interesting therapeutic option in diseases affecting the skin such as psoriasis or acnes vulgaris.

Here we have used mixtures of different skin microbiome components to alter the composition of a recipient skin microbiome.

**Results:** We show that after sequential applications of a donor microbiome, the recipient microbiome becomes similar to that of the donor. After intervention, an initial, week-long phase is characterized by dominance of donor strains. The level of engraftment depends on the composition of the recipient and donor microbiomes, and the applied bacterial load. We observed higher engraftment using a multi-strain donor solution with recipient skin rich in *Cutibacterium acnes* subtype H1 and *Leifsonia*.

**Conclusions:** We have demonstrated the use of living bacteria to modulate skin microbiome composition.

## Background

The human body is host to a complex and rich microbial community. The human microbiota mainly resides on the skin, oral mucosa, and in the gastrointestinal tracts, and has fundamental roles in health and disease^1^. The development of Next Generation Sequencing technologies has allowed the study of these communities with an unprecedented depth and resolution^2^. The gut microbiome has been investigated extensively^3^, with the skin microbiome becoming the focus of research more recently^4–8^. The skin is colonized by a large number of diverse microorganisms, of which most are beneficial or harmless^9^. More specifically microbes colonize the Stratum corneum of the epidermis and skin appendages such as sweat glands and hair follicles. The composition of abundant species is relatively stable over time^10^. However, skin-associated diseases such as acnes vulgaris^11^, eczema^10,12–14^, psoriasis^15^, rosacea^16^ or dandruff^17,18^ are associated with strong and specific microbiome alterations. For instance, the appearance of acne vulgaris has been linked to dysbiosis in the skin microbiome^11,19^. This distortion is probably caused by a specific subset of the skin bacterium *Cutibacterium acnes*^*11*,*19*–*21*^. Different strains of this bacterium have different degrees of association with acne. For instance, presence of strains carrying locus 2 is highly associated with the disease^21^. Conversely, different *C. acnes* strains have been associated with multiple positive properties^22^. The targeted manipulation of the human microbiome may become a potential therapeutic strategy for the treatment and study of diseases. The most prominent example of this therapeutic principle is the treatment of the antibiotic-resistant bacteria *Clostridium difficile* with the help of ‘fecal transplantation’^23^. Following this successful treatment a whole set of projects are developing microbiome-based treatments for gut diseases^24^. Similarly, manipulation of skin microbiome entails the promise of novel therapeutic approaches for skin diseases^25^.

We are particularly interested in *C. acnes* and its strain diversity, as this bacterium represents a major part of the human skin microbiome, and certain strains are associated with acne vulgaris^11,19,26^. Therefore, we developed and tested an approach to modulate the subpopulation of this species at the strain level.

## Results

In this work, we aimed to demonstrate that the human skin microbiome composition can be modulated through approaches similar to those used in fecal transplantation of the gut microbiome. For this, we prepared probiotic solutions from donor microbiomes, and applied them onto healthy volunteers, whose skin microbiome was monitored during and after the treatment. Two solutions comprise complete microbiome isolations from two donors (CM samples; CM1, CM2), and three others are composed of defined sets of *C. acnes* strains isolated from donors (PA solutions; H1, H1+A1,H1+D1+A1).

These solutions were applied on 18 healthy subjects with ages ranging from 22 to 42. Eight different skin areas were defined for application, whereof three were on the chest and five were located along the spine (Fig. 1A). These areas were chosen due to their typically high abundance of sebaceous glands. To get an understanding about the dose response of applied bacterial strains, three different cell concentrations were chosen (10^4^, 10^6^ and 10^8^ CFU/mL) and applied on the different areas. One area (area 4) was used as a negative control (i.e. no application). To better understand synergistic effects of strain combinations different strain combinations were used. One mixture contained only strain H1 (H1), a second was spiked with small amounts of A1 (H1+A1), and a third consisting of nearly equal amounts of H1, D1 and small amounts of A1 (H1+D1+A1). To circumvent biases on each subject area a different concentration was applied and rotated along the different individuals. All test areas except area 4 (control) were sterilized before application. Probiotic solutions were applied every day during days 1, 2, and 3. Skin microbiome samples were taken with commercial skin stripping method (3S-Biokit) based on fast hardening cyanoacrylate glue at 16 time points (0, 1, 2, 3, 4, 5, 8, 10, 12, 17, 24, 38, 52 days) to monitor microbiome dynamics (Fig. 1B). Genomic DNA was extracted and sequenced by NGS based genotyping. 16S rRNA gene profiling was used to assess microbiome composition at the genus level. SLST profiling^27^ was used to identify relative proportions of different *C. acnes* strains. Barcoded libraries were constructed and sequenced by an Illumina Miseq machine (Illumina, USA). The obtained data was quality filtered, mapped, and clustered (see Materials and Methods).

**Figure 1.**
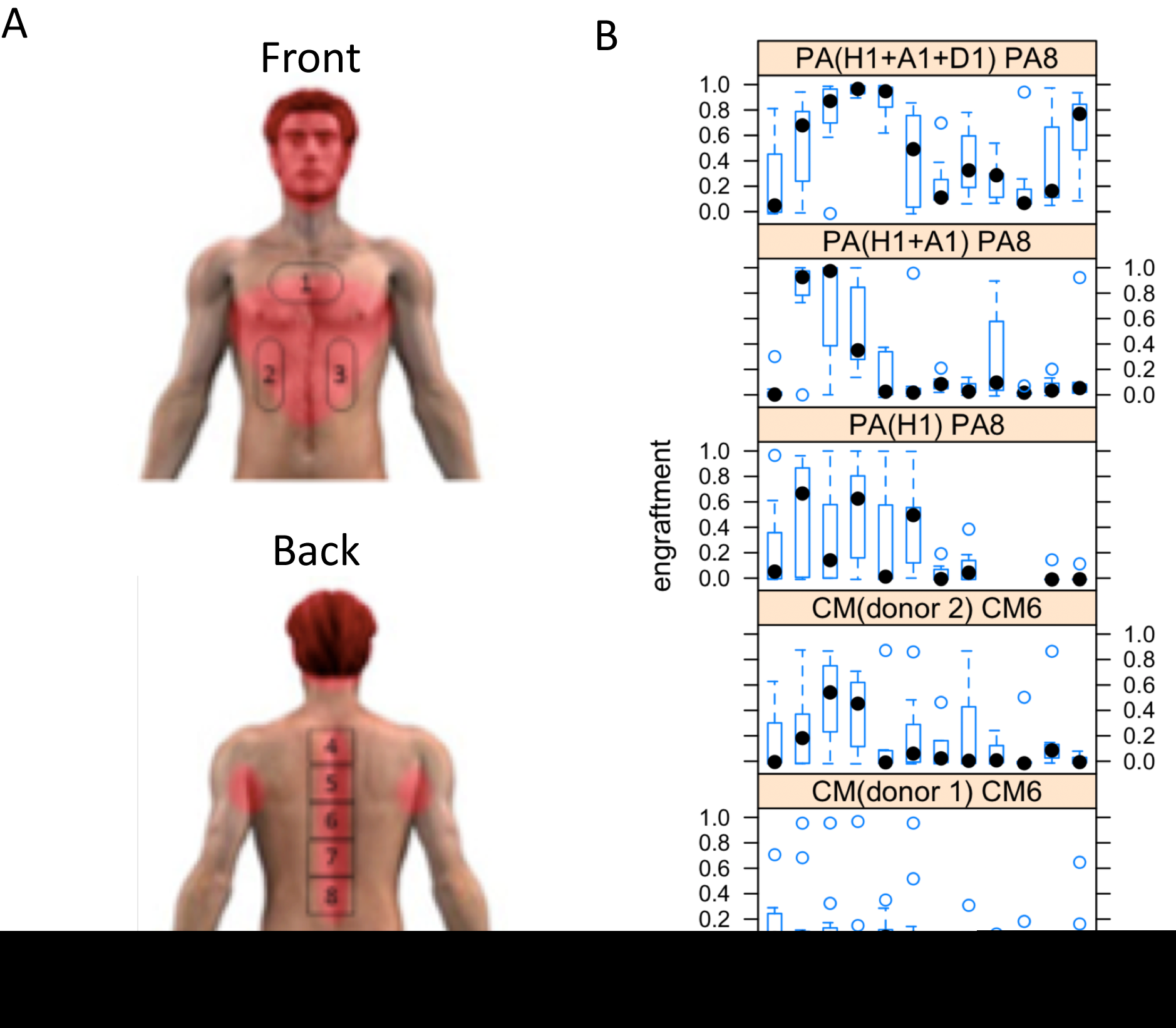
Skin microbiome composition dynamics after donor transplantation. A) Skin surface areas of study (in red: rich on sebaceous glands, squared: areas of application. B) Engraftment level of different probiotic solutions at different days of application

After the SLST profiling, we performed a partitioning around medoids (PAM) cluster analysis of the samples from all recipients at each time point based on Jensen-Shannon Divergence (JSD) distance (Fig. 1B), and used a Calinski-Harabasz (CH) index as well as average silhouette width to determine the optimal number of clusters^28,29^ (Fig. 2A). Based on this analysis we could identify five main clusters of skin population *C.acnes* profiles. We decided to name these five clusters dermatotypes 1, 2, 3, 4, and 5; analogous to the term ‘enterotype’ defined for the gut microbiome ^30^. The skin microbiomes of dermatotype 1 are driven by C. acnes L1; dermatotype 2 by C. acnes D1; dermatotype 3 by C3 and A5; dermatotype 4 by D1 and H1; and dermatotype 5 by C. acnes A1 (Fig. 2B). Second, we observed a quantitative and qualitative increase in similarity between donor and recipient microbiomes after only three days of application. For each solution, we assessed engraftment levels (Fig. 1B, 2C, Fig. S1), and the change of the composition of the *C. acnes* subpopulation before the treatment at three predetermined concentrations (10^4^, 10^6^ and 10^8^ CFU/mL). Engraftment is measured as the distance between the microbiome composition of the tested sample and the applied solution.

**Figure 2.**
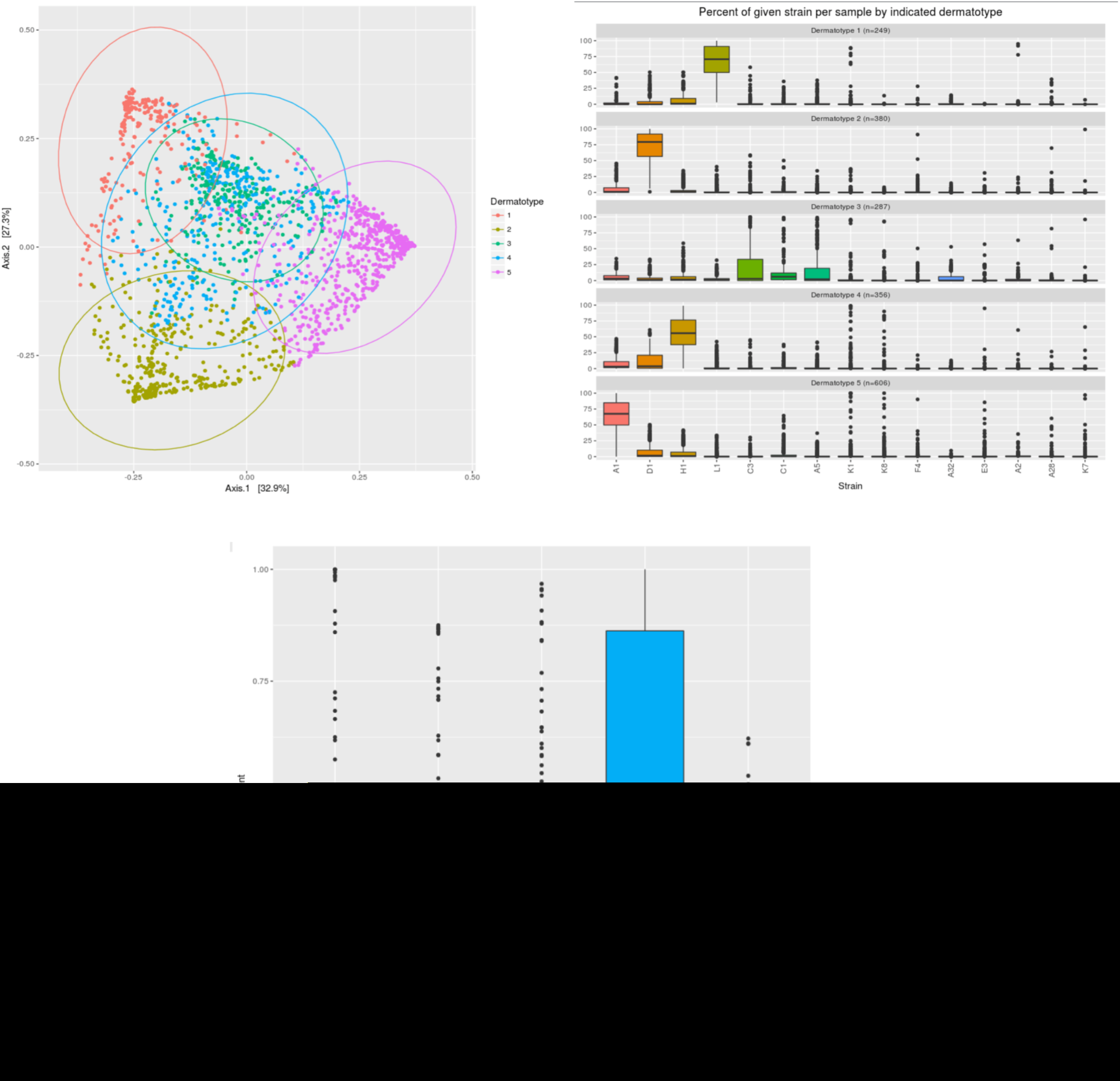
C. acnes population dynamics using SLST typing. A) PCA representation of the different dermatypes (based on SLST typing) B) Composition of the dermatypes (based on SLST typing) C) Average engraftment of different dermatypes (based on SLST typing)

Some of the applied mixtures engraft better. PA mixtures engraft better than CM at any concentration, and the highest concentration (PA8) has significantly higher engraftment values (Fig. S2). The values show that engraftment is greater with the (H1+A1+D1) solution, followed by (H1+A1) and (H1), in this order (Fig. S2). Expectedly, higher concentrations show greater engraftments (Fig. S2). PA8 containing H1, A1 and D1 engrafts significantly better than all the other groups.

Not all subjects responded equally to the applied samples, indicating significant variability among recipient areas that sometimes relate to defined *C. acnes* based dermatotypes. For instance, dermatotype 4, shows higher engraftment than others (Fig. 2C, Tukey test). Interestingly, this dermatotype is dominated by H1, and comprises notable levels of D1 and A1 (Fig. 2B).

We also classified patients according the different 16S based dermatotypes. In this case we observe 3 different types: type one dominated by Cutibacterium, type two dominated with Cutibacterium and some Corynebacterium, and a more widespread type 3, with Leifsonia being the most abundant (Fig. 3A, Fig. 3B). Patients with type 3, show a significantly higher engraftment (Fig. 3C). We hypothesize that patients of type 3 are not fully colonized with *Cutibacterium* and therefore it is easier to establish a new population.

**Figure 3.**
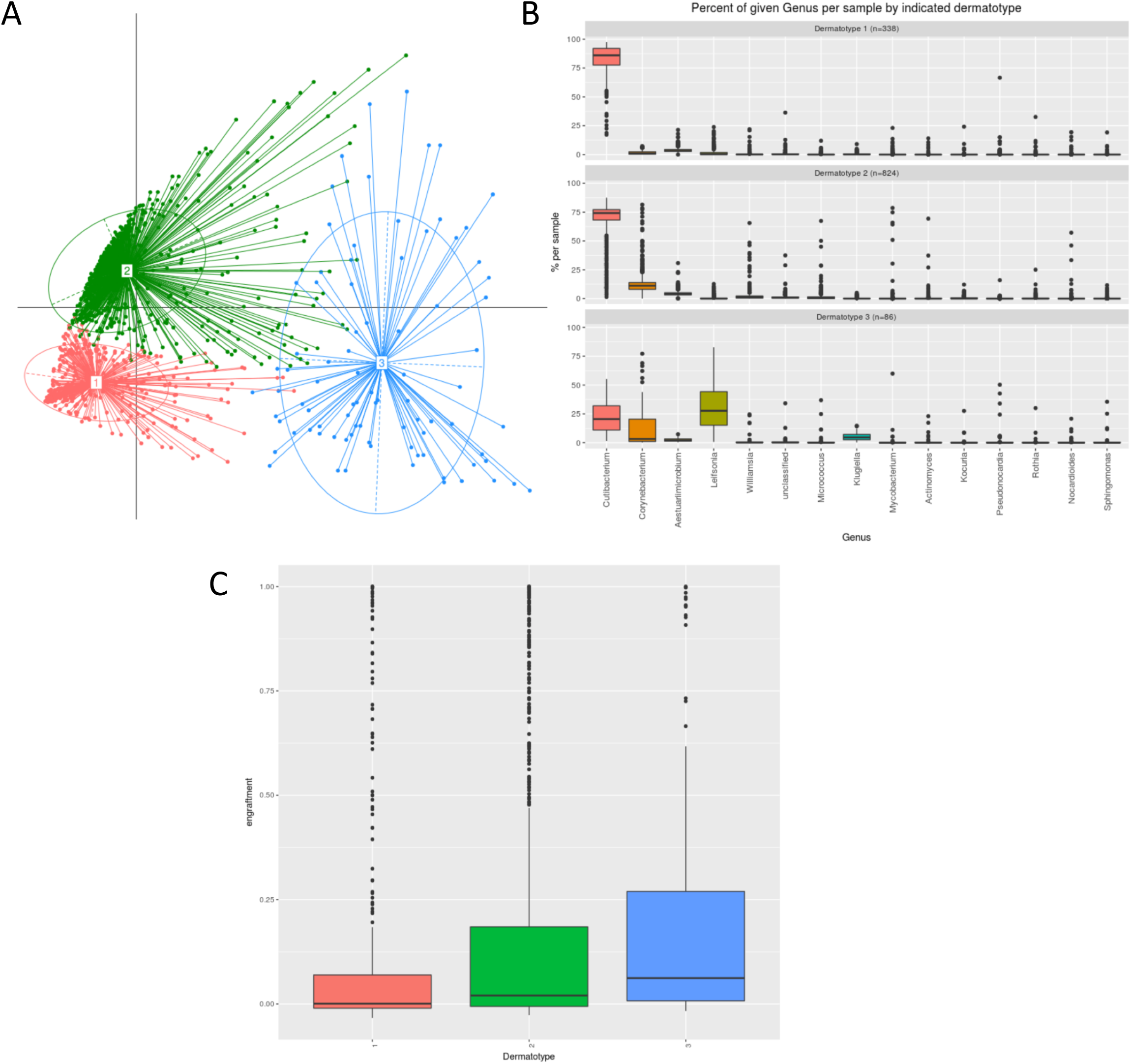
Complete microbiome dynamics at the 16S level. A) PCA representation of the different dermatypes (based on 16S typing) B) Composition of the dermatypes (based on 16S typing) C) Average engraftment of different dermatypes (based on 16S typing)

## Discussion

We have demonstrated that the composition of the human skin can be modulated by applying *C. acnes* strain H1 with positive features isolated from healthy individuals^31–33^ The dose of applied bacteria plays an important role in the modulation capacity.. During the first three days, the abundance of applied bacteria increases with every day and decreases gradually after termination of application. The applied dose determines the prolongation of the abundance of the applied strain on the tested skin.

This return to the ground state is in accordance with a recent study reporting temporal stability of the skin microbiome^5^. Unfortunately our data is limited to skin areas rich in sebaceous glands and we do not know whether other areas which are suspected to be more dynamic^6^ and involved in other diseases^13^ react differently. This is an interesting question as different body sites harbor different *bacterial* subpopulations^27^ and their reaction to external modulation might be different.

## Conclusions

Microbes are important components of the skin. Recent clinical studies already revealed that application of natural bacteria into the skin can decrease skin pH and improve moisture retention^34^. This method opens the possibility to develop probiotic solutions that help the human skin reverting disease microbiome states to healthy ones. Also, synthetic biology is generating smart microbes with the abilities to detect and treat disease^35^. New methods to replace, and modulate our bacterial flora are extremely necessary. We expect that this methodology could be used to study and modify skin microbial components and have broad implications for future therapies and research in skin microbiome and related diseases.

## Methods

### Isolation from the donor

A mixture of biologically active probiotic bacteria for topical administration was prepared as follows. A sample of skin microbiome was taken from a donor. The sample was then cultured in the laboratory, and a formulation was prepared.

Methods for analyzing the microbiome included DNA isolation, 16S amplification and large-scale amplicon sequencing, as well as bioinformatics for the taxonomic assignment and quantification of diversity in microbial communities. Steps included:

1. Isolation of bacterial strains from a donor. Bacteria were collected using swabs.
2. Growth in the laboratory. Bacteria were grown in reinforced clostridium agar in anaerobic conditions at 37°C.
3. Isolation and manipulation of the bacterial strains. The sample was enriched for 20 *Cutibacterium* strains and analyzed for positive genotypes with SLST primers.
4. Formulation of a probiotic. Bacteria were collected from an agar plate using a saline solution.
5. Application of the probiotic to the recipient. The donor microbiome was applied using swabs.
6. Genotyping of the modified recipient microbiome, using an NGS-based genotyping approach discussed below.

### Skin microbiome donor preparation viability

The *C. acnes* strain 6609 was grown in Reinforced clostridal medium as liquid culture. After 2 days, the culture was spun down and washed with PBS (Phosphate buffered saline, pH 7.4) and as a final wash with water. Then the culture was resuspended in pure water and aliquoted into samples. A concentrated solution of the additives tested 10× PBS was added to the bacterial suspension. Then the aliquots were stored either at room temperature or at 4°C. In both cases, they were protected from Sunlight. In regular intervals, about every 3-4 days, a dilution series of each sample was taken and the colony-forming unit (CFU) count was determined. The suspension was vortexed and a serial dilution was prepared. To determine the CFU count, aliquots of the dilution were added on a agar plate which is suitable to grow C. acnes. 10μl of an appropriate dilution was added on an agar 30 plate of reinforced clostridial medium. The 10μl was placed as a drop on top of the plate and run down. This method allows the placement of up to 4 drops on the plate. (e.g., see http://www.science-projects.com/serdil.htm). Each sample was determined in 4 technical replicates. After 3-4 days of anaerobic incubation, the colony numbers were counted (manually or using the software OpenCFU) and both the average and the standard deviation were determined. Thereby a profile of the colony forming units was monitored over time. Additionally, bacteria of the skin microbiome were stabilized in a neutral liquid matrix for several days at room temperature. It was demonstrated that *Cutibacterium* can survive weeks of storage at room temperature. Constant numbers of colony forming units (CFU) from a liquid matrix over a week were also recovered. To assess these numbers, methods that determine the CFU of liquids in a medium-throughput fashion were established as described below. It was shown that compositions were stable for at least 1.5 months. Longer time periods are being tested.

### Donor microbiome solution application to the recipient

Microbiome donor solution was applied once a day during 4 days using swaps in to a delimited area on the chest of the recipient (Fig. 1S). Prior to application the area was cleaned and disinfected. Sampling for genotyping was carried out before new donor samples were applied.

### Strain genotyping

An NGS-based genotyping approach was used for identifying different strains:

1. The microbiome was collected using swabs daily
2. The sample was incubated at high temperature to isolate the DNA. The QuickExtract^™^ kit from Epicentre, Chicago, IL, was used with some modifications. 80 microliters of 0.05M NaOH was added to the suspension solution. The incubation was conducted for 45 minutes at 60°C, followed by a 5 minute incubation at 95°C. After incubation, 920 microliters of 1 M Tris-HCl, pH 7.0 was added. 0.5 microliters of this mixture was used for PCR.
3. PCR was conducted on the sample using 16S primers, and SLST allele to characterize the population. DNA preparation was diluted 100 × for PCR analysis. Samples were amplified using KAPA polymerase (5 min 95°C; 35 cycles of (98°C 20s, 62°C 25s, 72°C 30s); 1 min 72°C
4. Library preparation. The library was constructed using two rounds of PCR. The 10 first round used 16S primers and SLST primers which included sequences compatible with Illumina sequencing. The second round was used to barcode the different samples for sequencing in a single Illumina flowcell.
5. Illumina MiSeq sequencing was conducted.
6. Samples were analyzed using an internally developed computational pipeline (S-genotyping). Quality filtering; samples were mapped into an internal database using bwa software; data processing and visualization was conducted with R statistical language.

### Normalization and filtering of 16S and SLST data

The 16S rRNA gene counts and the SLST counts for the samples in this study were stored and analyzed using the R package Phyloseq (version 1.16.2)^36^. The counts were normalized per sample by dividing each value by the sum of all counts for a given sample and multiplying by 100, leaving the relative abundance of each genus/strain within that sample, with all values between 0 and 100.

### Clustering and dermatotype analyses

The Jensen-Shannon Divergence (JSD) was used to produce a distance matrix between the genera/strains of all samples and then partitioning around medoids (PAM) clustering to group samples with similar overall abundances. We used the Calinski-Harabasz (CH) index to determine the optimal number of clusters, and we further verified this by calculating the average silhouette width of the samples, which is a measure of the separation of samples within one cluster from those of another cluster. The functions for these calculations come from the R packages cluster (version 2.0.4)^37^ and clusterSim (version 0.44-2)^38^. A Principal Coordinate Analysis (PcoA) was used to visualize the clustering of the samples within their respective dermatotypes with the R package ade4 (version 1.7-4)^39^.

## Declarations

### Ethics approval and consent to participate

All procedures in this work were carried out following the principles expressed in the Declaration of Helsinki and have been approved by Universitatkilinikum Magdeburg IRB (ref. 171/15 – Dose-response in vivo testing of direct manipulation of the skin microbiome using natural bacteria).

### Availability of data and material

Genomics datasets have been submitted to EBI ENU with project ID PRJEB28732

## Acknowledgments

We thank María Angélica Martínez for advice and help on initial stages of the project.

## Author contributions

BP, and MG designed and performed the experiments. BP expanded the microbiome cultures ex vivo. SRQ designed and supervised the microbiome application to skin and collection from skin. TG and MG performed the computational analysis. BP, MG, and TG wrote the manuscript.

